# A new computational model captures fundamental architectural features of diverse biological networks

**DOI:** 10.1101/046813

**Authors:** Bader Al-Anzi, Noah Olsman, Christopher Ormerod, Sherif Gerges, Georgios Piliouras, John Ormerod, Kai Zinn

**Affiliations:** Division of Biology and Biological Engineering; Control and Dynamical Systems Option, Division of Engineering and Applied Sciences; Division of Physics, Mathematics, and Astronomy, California Institute of Technology, Pasadena, CA 91125; Lewis-Sigler Institute for Integrative Genomics, Princeton University, Princeton, NJ 08540; Singapore University of Technology and Design, Engineering Systems and Design (ESD), 8 Somapah Road, Singapore 487372; School of Mathematics and Statistics F07, University of Sydney NSW 2006, Australia

## Abstract

Complex biological systems are often represented by network graphs; however, their structural features are not adequately captured by existing computational graph models, perhaps because the datasets used to assemble them are incomplete and contain elements that lack shared functions. Here, we analyze three large, near-complete networks that produce specific cellular or behavioral outputs: a molecular yeast mitochondrial regulatory protein network, and two anatomical networks of very different scale, the mouse brain mesoscale connectivity network, and the *C. elegans* neuronal network. Surprisingly, these networks share similar characteristics. All consist of large communities composed of modules with general functions, and topologically distinct subnetworks spanning modular boundaries responsible for their more specific phenotypical outputs. We created a new model, SBM-PS, which generates networks by combining communities, followed by adjustment of connections by a ‘path selection’ mechanism. This model captures fundamental architectural features that are common to the three networks.

## Introduction

Essential insights into basic cellular processes have been obtained through the biochemical and genetic analysis of multimeric protein complexes that perform specific functions. In an analogous manner, an understanding of the functions of small regions or circuits in neural systems has been attained through detailed analysis of synaptic communication among individual neurons or groups of neurons. Cells contain large numbers of interacting molecular complexes, while neuronal circuitries are composed of interconnected neurons or groups of neurons. These elements are organized into highly connected communities, typically referred to as modules, which themselves are connected to one another to form complex networks capable of generating ‘phenotypic outputs’. Cellular phenotypic outputs include processes such as cell division, while phenotypic outputs for neural systems can be specific behaviors (*e.g.*, courtship) or physiological responses (*e.g.*, heart rate regulation). However, the phenotypic outputs of a given protein or neuronal structure are not necessarily predicted by the general function of the module to which they belong. For example, cell morphology as a phenotypic output requires the cooperation of proteins that include components of transcription complexes, translational regulators, protein-modifying enzymes, and the cytoskeleton. An understanding of how these modules function as separate entities does not necessarily define the phenotypic outputs in which they participate.

Each module within a cell or a nervous system can potentially contribute to many different phenotypic outputs, each of which might utilize different components within participating modules. However, the relationships linking interactions between components within a module (proximal interactions) to the interactions of those components with members of other modules (distal interactions) responsible for controlling a phenotypic output are not well understood.

In order to better understand large biological networks containing hundreds of elements, it will be necessary to move beyond the experimental analysis of the functions of individual components by using computational models. These models can simulate the architectures of entire networks and analyze how network structure influences its functions. Computational models can predict how deletion or modification of individual elements within a network will affect its overall architecture, and can thus identify key elements within a network that are likely to be essential to its functions and can become targets for experimental analysis.

Biological networks are often analyzed using graph theory. In graph-theoretical models, genes or proteins are represented by nodes/vertices, while their corresponding transcriptional or proteomic interactions are represented by edges. Similarly, in graphs corresponding to neural circuitry, neurons or small groups of neurons in the same neuronal structure would be represented by nodes/vertices, while the axonal tracts connecting them would be represented by edges. Central to network analyses are measures reflecting their connectivity and physical topology(1). These include: 1) the degree distribution *P(k)*, which is the probability that a randomly chosen vertex has degree *k*; 2) the clustering coefficient *Cg*, which measures how often collections of nodes form small highly interconnected groups; 3) the mean path length *L*, which is the average number of steps along the shortest paths that connect all possible pairs of network nodes; 4) the modularity *M*, which measures the division of a network into modules as result of the tendency of some nodes to form connections primarily within the module to which they belong. Collectively, these terms encompass the fundamental properties of network architecture and are frequently used to describe real-world networks(2, 3),(4, 5).

It has been widely believed that most biological networks are ‘scale-free’, because they are often characterized by a *P(k)* distribution that resembles a power law, due to the presence of select ‘hub’ nodes that maintain exponentially more connections than other nodes within the network. They also have high *Cg* and *M* values along with small-world properties that allow nodes to reach each other via a small number of steps(6, 7).

The four computational models ferquntly used to analyze biological networks are the Erdos-Renyi (ER)(8), Barabási-Albert (BA)(9), Watts-Strogatz (WS)(10) and Hierarchical random graph (HRG) models(11). The ER model is characterized by an equal probability of forming connections between any two nodes. The BA model generates graphs using a preferential attachment process. In this mechanism, more highly connected nodes are more likely to receive new links and thus become hubs. Networks generated by the BA model are scale-free, i.e., they have *P(k)* distributions that are described by a power-law. In the WS model, networks are generated by randomly rewiring a regular ring lattice. This algorithm was designed to produce better models of real-world networks by remedying the lack of clustering seen in ER and BA networks. Finally, the HRG model is based on the replication of the hierarchical organization of nodes into modules and submodules. The probability of forming an edge between any two nodes depends on how closely the two nodes are related in the hierarchy.

The use of these computational models has not yet provided major insights into the functions of real networks, for a variety of reasons. First, the networks chosen for analysis are often generated by sampling methodologies that assemble elements without regard to common physiological function. For example, proteomic methods can identify all connected proteins in a cell via genome-wide techniques. Some investigators have analyzed the entire network of cellular proteins, while others have examined highly connected essential proteins whose removal produces a lethal phenotype(12-19). These approaches do not account for the fact that only select nodes and edges are relevant to specific cellular functions, that proteins may use different partners to regulate different phenotypic outputs, and that lethality does not necessarily imply a common biological function. Second, real biological networks have major features that significantly differ from those predicted by the models described above. For example, ER models produce networks with low *Cg* values, and the WS model produces networks with binomial/Poisson *P(k)* distributions that rarely encountered in biological networks. Also, WS networks are generated from an initial highly ordered ring lattice, and this is not a biologically plausible mechanism for network evolution. The BA model produces networks with a *P(k)* distributions that follow a power-law(20), but these networks have low *Cg* values(20). Third, the dogma that biological networks have power-law *P(k)* distributions may not hold up to rigorous statistical scrutiny, since combinations of Poisson/binomial-like degree distributions can produce distributions that superficially resemble a power-law. Indeed, when published biological networks were subjected to detailed mathematical analysis in which a variety of *P(k)* distributions types were tested, including Poisson, binomial, and power-law, none of the models could be definitively proven or ruled out(21, 22). Network models such BA have focused on the power-law *P(k)* distribution aspect of biological networks above other features. However, illustrated in Supplementary Figure 1, this feature in isolation from other topological elements cannot be diagnostic for a given network type. Finally, these models do not explain the variety of phenotypic outputs generated by real-world networks.

In our earlier work, we attempted to address these problems by experimentally identifying ~100 genes required for fat storage control in budding yeast(23). We showed that proteins encoded by these genes define a highly interconnected network. By definition, proteins in this network share at least one common function, that of fat storage regulation. We thus we avoided the problems encountered by global protein networks generated from protein-protein connections that do not reflect common biological functions. We found that, unlike other published biological networks, the fat storage regulation network is not scale-free and could be simulated using the WS model. We also showed that the value of a topological parameter called Katz centrality correlated with the importance of a given protein to network function, and we used the network architecture to design pharmacogenetic experiments that defined its patterns of signal flow(23).

In the work described in this paper, we applied the approaches we had used for the fat regulatoion network to several larger networks. In doing this, we also hoped to see if the methods we had developed for analysis of proteomic networks could be applied to anatomical networks. In selecting appropriate networks for analysis, we employed two major criteria. First, the nodes of the networks should be linked by common functions. Second, the networks should be as complete as possible, so that they are unlikely to be missing large numbers of nodes or edges. This criterion ensures that a computational model of the experimentally derived network is likely to reflect the actual network that exists *in vivo*.

We found three near-complete large biological networks fulfilling our criteria: 1) the network of all proteins required for mitochondrial function in budding yeast; 2) the mesoscale connectome of the mouse brain; and, 3) the complete neural network of the *C. elegans* hermaphrodite. These three networks are similar only in that they are all products of evolution and all generate phenotypic outputs (cellular physiology for the mitochondrial network, and animal behavior for the mouse and worm networks). We reasoned that a model capturing essential features of all three networks would provide evidence that we uncovered fundamental properties shared by real biological systems.

The results reported here show that all three networks are composed of large communities containing modules with general functions, containing topologically distinct subnetworks spanning modular boundaries responsible for specific phenotypic outputs. Their common architectural features led us to develop a new and evolutionary plausible network model we denote as the Stochastic Block Model with Path Selection (SBM-PS). In this model, networks are generated by linking different communities (blocks) *via* connections that form with higher probability within a community than outside of the community. This is followed by removal of connections with probabilities dependent on their contribution to short paths of communication. We show that the SBM-PS captures the essential architectural features of all three networks with higher probability than any other currently available computational model, and that it produces *Cg* and *M* values that are prerequisites for the formation of subnetworks that execute phenotypic outputs.

## Results and Discussion

### Choice of networks for analysis

We refer to the three networks analyzed in this study as the yeast mitochondrial network, the mouse brain mesoscale connectome, and the *C. elegans* connectome. We generated the yeast mitochondrial network using bioinformatics, first identifying all yeast (*Saccharomyces cerevisiae*) genes for which loss of function (LOF) had been shown to produce abnormal or dysfunctional mitochondria in published studies. We then used global proteomic data to generate a network representing all molecular interactions among the proteins encoded by these 883 genes(12-17, 23-27) (Figure 1A, Supp. Table 1). Remarkably, close to 85% of the genes encode proteins that are physically connected, forming an extensive network. The majority of the connections in the network (~80%) were identified by co-precipitation methods that detect stable and abundant protein complexes that exist *in vivo*(14, 15). Given that mutations eliminating network proteins ultimately affect mitochondrial function (*i.e.*, they share a set of phenotypes), the connections in this network are relevant to cellular processes affecting mitochondrial structure and physiology. Because the genes were identified in multiple genetic screens of the entire yeast genome, including essential genes(26), the network is likely to include all proteins necessary for mitochondrial function. However, it would lack proteins encoded by redundant genes. We also evaluated the biological significance of the connections in the network by examining the ability of randomly selected sets of equal numbers of proteins from the yeast genome to form a network *via* their annotated proteomic interactions. All such networks had less than half as many edges as the mitochondrial network, and they all had low *Cg* values (Supplementary Figure 2). These data indicate that the interconnectedness of the yeast mitochondrial network was selected by evolution.

**Figure 1.**
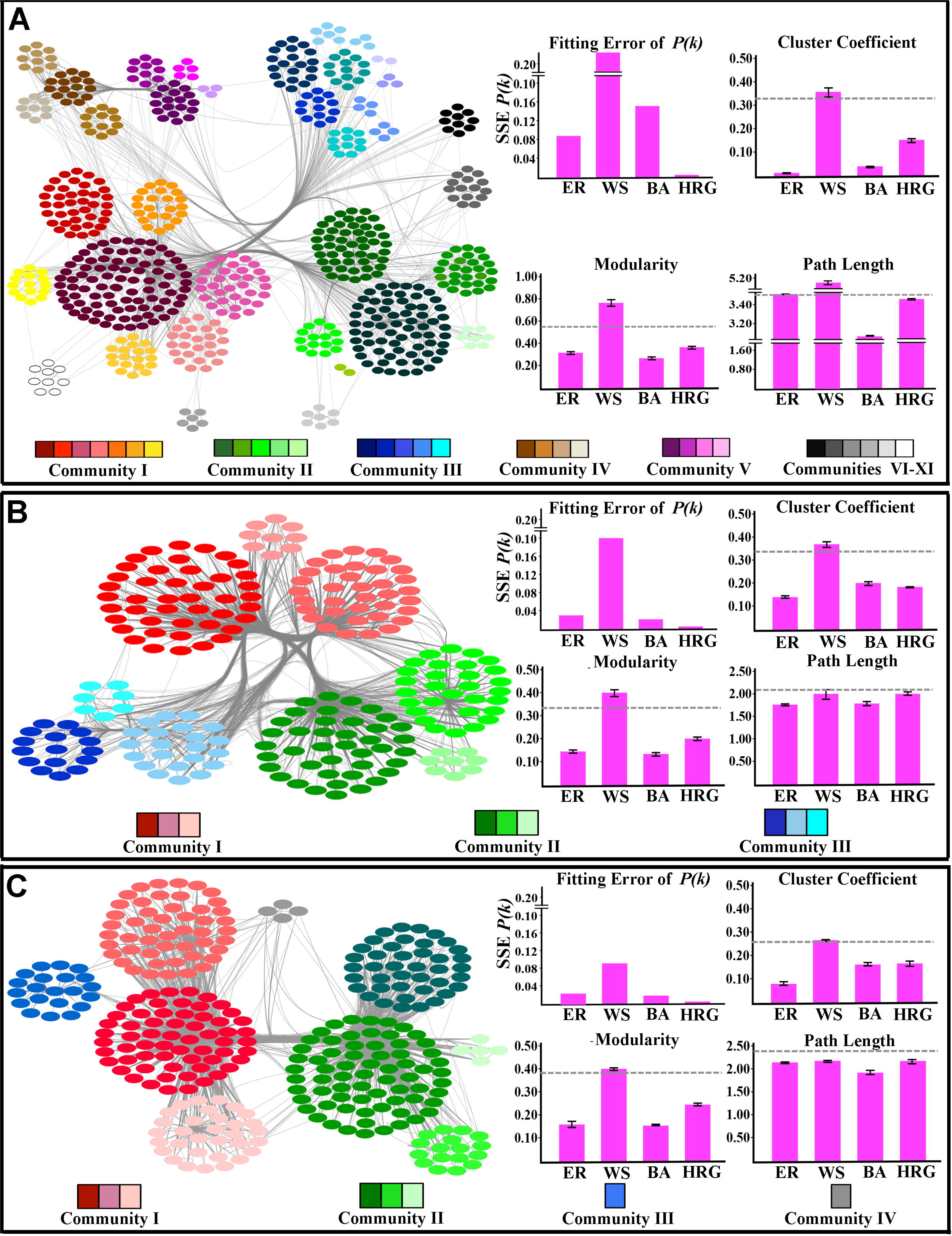
Community structures of the yeast mitochondrial network (A), mesoscale connectome of the mouse brain (B), and *C. elegans* connectome (C). These networks are composed of large communities that contain smaller sub-communities. Community and sub-community boundaries were delineated using the walk-trap algorithm. Connections between nodes (edges) are shown by light gray lines. Nodes that belong to the same large community are represented by different shades of a particular color (see color bars at bottom of each panel), while nodes that belong to the same sub-community have the same shade. The identities of nodes (corresponding to proteins in the yeast mitochondrial network, brain regions in the mouse brain, and single neurons in *C. elegans*) are shown in Supp. Fig. 2. The histograms on the right sides of each panel compare each real network to collections of 20,000 simulated networks captured by four computational models (ER: Erdos-Renyi; BA: Barabasi-Albert; WS: Watts-Strogatz: Hierarchical model; HRG). The *P(k)* histograms show the fitting error, as measured by SSE, between the *P(k)* distributions of the real networks and the simulated networks. The *Cg*, *M*, and *L* histograms show the averages of these topological parameters for the collections of simulated networks. The grey dashed line in each histogram indicates the value of this parameter in the real network. Error bars indicate standard deviations.

The mouse brain mesoscale connectome network was produced by The Allen Institute for Brain Science by injecting a recombinant adeno-associated virus (AAV) expressing EGFP as an axonal anterograde tracer into the mouse brain(4) (Figure 1B, Supp. Table 2). The tracer was applied to 295 non-overlapping anatomical regions that span most of the mouse brain(4). Only 18 areas in the brain were not labeled due to problems with tracer injections.

Finally, the adult *Caenorhabditis elegans* hermaphrodite contains 302 neurons, and the axonal connections among these neurons were completely defined using electron microscopic (EM) reconstruction(28-30) (Figure 1C, Supp. Table 3).

We did not attempt to examine gene regulatory networks in this analysis. These are composed of genes and regulatory proteins that interact with them, which may also be products of genes within the network. Unlike proteomic networks, the relationships between the elements forming them and the physiological responses they regulate are not necessarily direct. For example, transcribed mRNAs may not be translated into protein, and not all transcripts are necessarily involved in the physiological response of interest. By contrast, the yeast mitochondrial network and other protein networks are derived from the final products of gene transcription. For the mitochondrial network, we also know that all of its components are involved in the same physiological process, since mutations eliminating them ultimately affect mitochondrial function.

### The yeast mitochondrial network and the mouse brain and *C. elegans* connectomes have similar topologies

To analyze the three networks, we first calculated their topological parameters and then evaluated whether similarly sized networks with similar parameter values could be generated using the four common algorithms discussed above. All three networks have high *Cg* and *M* values and short path lengths (*L*) consistent with ‘small-world’ properties (Figure 1). These values indicate that the networks contain densely interconnected modules, with sparser connections between nodes that belong to different modules.

Unlike the two neural networks, which are composed from edges defined using the same methodologies (tracer injection for the mouse and EM reconstruction for the worm), close to 20% of the edges in the mitochondrial network were not detected by co-precipitation methods, but using yeast two-hybrid or biochemical phosphorylation assays, raising the possibility that these edges are qualitatively different from the majority of edges in the network and should not be grouped with them in a topological analysis. We thus recalculated *Cg*, *L*, and *M* for the network after removal of these edges, and we did not observe any dramatic changes. *Cg* changed from 0.33 to 0.38, while *M* changed from 0.58 to 0.65, and *L* from 3.62 to 3.86.

We generated 20,000 simulated networks with equivalent numbers of nodes and edges to the three experimental networks using the ER, BA, WS, and HRG algorithms, and computed the empirical distribution of the topological parameters of networks generated by each model. When we compared the *P(k)* distributions of the real networks with the BA model, representing a best-fit power-law distribution, or the ER model, representing a best-fit Poisson/binomial distribution, we found that there were substantial fitting errors, as measured by the sum of squares due to error (SSE) (Figure 1). This is particularly obvious for the yeast mitochondrial network (Figure 1A). These results indicate that the *P(k)* distributions of the real networks do not follow either a power-law or a Poisson/binomial distribution.

When we examined the other key topological parameters, we found that the BA, ER, and HRG models generate *Cg* and *M* values that are much lower than those of the real networks (Figure 1). The WS model can be chosen to reproduce a given *Cg* value and produce high *M* values, but it has the worst fit with respect to the *P(k)* distributions out of the tested network models. Thus, none of the four algorithms can generate the combination of topological features observed for the three real-world biological networks.

### The three networks are divisible into communities

In order to analyze the architectures of the yeast mitochondrial, mouse brain and *C. elegans* networks, we partitioned them into communities using a walk-trap algorithm (Newman Fast Community Finding Algorithm), considered to be a canonical method for community detection(31). This method is based on the assumption that short random walks from one node to another will tend to stay in the same community. One of the shortcomings of this approach is that the behavior of the walk-trap algorithm can be sensitive to small changes in the network. That is to say that the removal of single nodes could potentially alter the community assignment for a large number of nodes (for example, missing a node or an edge). We addressed this by subjecting the community structure obtained to a stability analysis. Our results indicate that nearly all nodes in all three networks stay in a given community under small changes in the network, indicating a relatively high degree of confidence that all our nodes belong in their assigned community (Supp. Tables 4, sheet 1, sheet 2, and sheet 3).

Our analysis indicates that the yeast mitochondrial network is composed of five large communities (each with >20 nodes) and seven smaller ones (Figure 1A, Supp. Table 1, sheet 1). The mouse brain mesoscale connectome is divided into three communities (Figure 1B, Supp. Table 2, sheet 1). Finally, the *C. elegans* connectome contains three large communities (each with more than 20 nodes) and one small community (four nodes) (Figure 1C, Supp. Table 3, sheet 1). (Supplementary Figure 3 shows the same diagrams, but with the names of components included.) By repeating the analysis performed for the entire graph on each isolated community, we find that the *P(k)* distributions of the large communities resemble Poisson/binomial distributions (Supp. Table 5).

Interestingly, the properties of the large communities resemble those of the fat storage regulation network we characterized in our earlier work (23), which is similar in size (~100 proteins) to the larger communities in the yeast mitochondrial network. The fat storage regulation network also has a Poisson/binomial-like *P(k)* distribution, and it can be accurately modeled using the WS algorithm(23). Fat storage regulation requires mitochondrial function, and most of the proteins in the fat storage regulation network are also members of the yeast mitochondrial network. Thus, much of the fat storage regulation network can be considered to be a subset of the mitochondrial network.

In examining the networks, we noted that large communities appeared to be preferentially made from nodes corresponding to proteins, areas, or neurons that are proximal to each other within the cell or nervous system. To evaluate this hypothesis, we assigned each node to a given cellular or neuronal compartment based on previously published literature (30, 32-34), and these results are depicted in the diagrams of Figure 2. For example, in the case of the yeast mitochondrial network, almost all nodes (proteins) in communities II, IV, and V are located in the mitochondrial compartment, while most nodes in community III are located in the nucleus (Figure 2A). In the *C. elegans* connectome, almost all nodes (neuronal cell bodies) in community I are in head ganglia, while most nodes in community II are in tail ganglia and the ventral nerve cord (Figure 2C). In the mouse brain mesoscale connectome, most community II nodes (brain areas) are in the limbic system. Community I is split between the isocortex and the brain stem, while community III is split between the cerebellum and brain stem (Figure 2B; Supp. Table 3, sheet 2). (Supplementary Figure 4 shows the same diagrams, but with the names of components included.) To determine if the tendency of nodes within the same large community to be located in the same compartment is statistically significant, we calculated the probabilities that the observed distributions of nodes among compartments in the three networks could be generated by chance. Out of 21 discrete structural groups, 17 had *p*-values based on a chi-squared test less than 10^−3^, implying that the community structure is statistically significant. Therefore, it is likely the walk-trap algorithm groups’ nodes that are close spatially into the same community.

**Figure 2.**
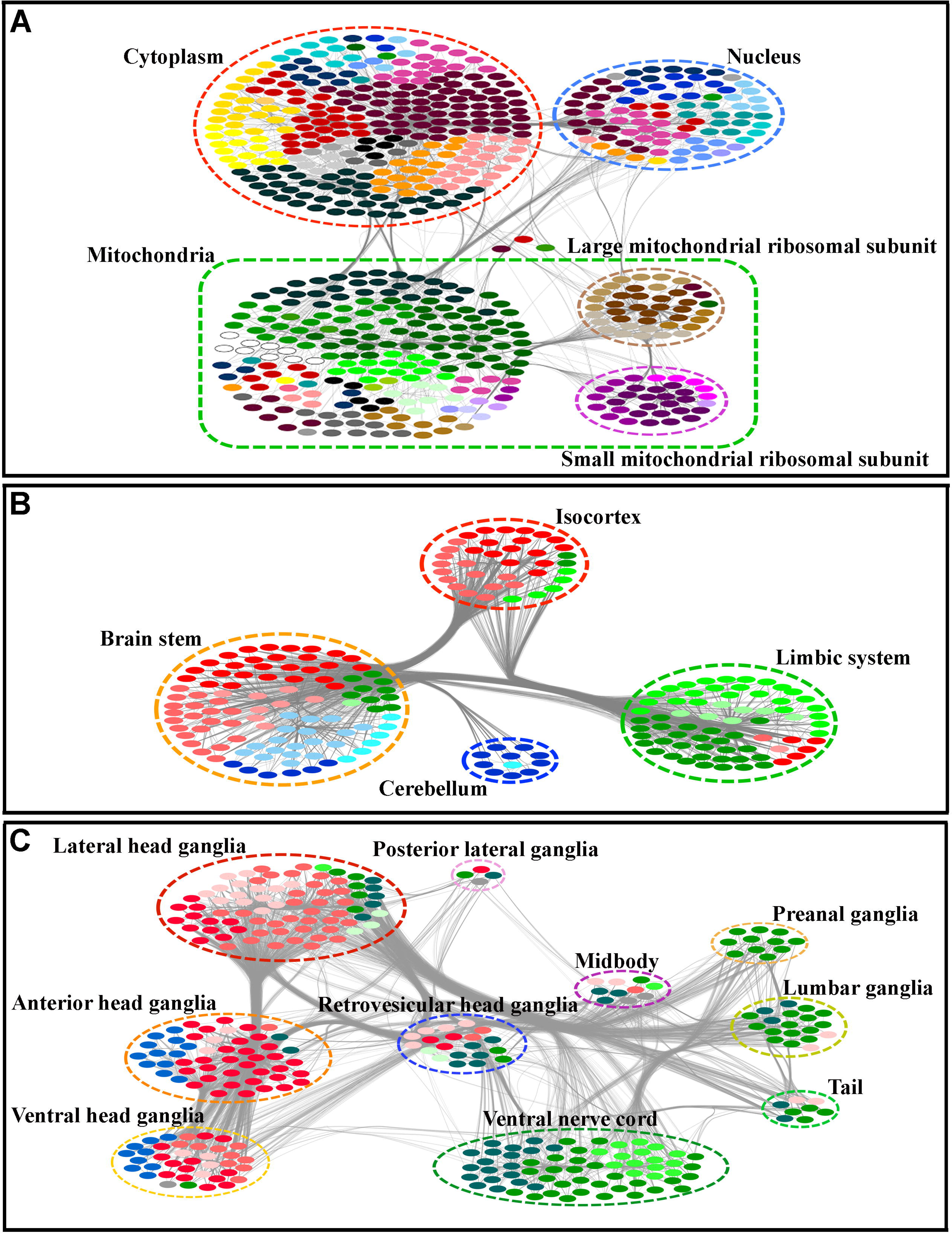
Spatial localization of network nodes. In the yeast mitochondrial network (A), the locations of nodes (proteins) within the cell are indicated by the labeled dotted outlines. The mitochondrial ribosomal subunit communities are indicated as enclosed within the mitochondrial outline. In the mesoscale connectome of the mouse brain (B), the locations of nodes (brain regions) within four large brain subdivisions are indicated by the labeled dotted outlines. In the *C. elegans* connectome (C), the locations of nodes (neuronal cell bodies) within ganglia are shown. The color code indicating community and sub-community identity is the same as in Figure 1. Light gray lines delineate edges. Identities of nodes are indicated in Supp. Figure 3.

The large communities in the mouse brain network perform diverse biological tasks and do not belong to a single functional category. Similarly, communities I, II, and III in the yeast mitochondrial network span functional categories. However, communities IV and V correspond to the large and small mitochondrial ribosomal subunit proteins, respectively, so they do have unified functions. For the *C. elegans* connectome, each large community has multiple functions. However, community I has many neurons with sensory functions, while community II is enriched in neurons with motor functions.

### Division of networks into sub-communities and phenotypic subnetworks

When each of the large communities in the three networks were separated from the rest of the network and subjected to further walk-trap analysis, we noted that they consist of smaller sub-communities composed of nodes that are even more proximal to one another either as result of being in the same molecular complex (in the case of the yeast mitochondrial network), the same brain nuclei (in the case of the mouse brain network), or the same ganglia (in the case of the *C. elegans* connectome) (Supp. Tables 1, 2, and 3, sheet 1). Furthermore, unlike the larger communities, these smaller sub-communities, which we denote as modules, often exhibit unified functions that can be utilized by different biological processes. One example is the mitochondrial sub-community I, G. which is largely composed of the SCF ubiquitin-protein ligase complex and its substrates(35) known to be involved in glucose detection and cell cycle regulation^21^. Another example is the mouse brain sub-community II, B, which is composed of amygdalar structures that are required for memory, decision making, and emotional reactions(33). Finally, worm connectome sub-community I, B is composed of the amphid chemosensory neurons, which are utilized in aggregation/dispersion behavior and chemosensory responses(30).

The phenotypic outputs of nodes within protein networks can be evaluated through examination of the phenotypes that are generated when the node is removed by knocking out the gene that encodes it. The phenotypic output of nodes within neural networks can be examined by lesioning neurons or altering their activity. For mouse brain nodes, this would be done by surgical or pharmacological lesions of a given area, or by using viral vectors to alter neural activity of neurons within a given area. The *C. elegans* connectome nodes correspond to single neurons. These can be laser-ablated, or their activity can be selectively altered using transgenic methods. We assembled collections of nodes involved in various phenotypic outputs from the three networks in an unbiased manner by listing all yeast mitochondrial network proteins, mouse brain areas, and worm neurons that had been associated with those outputs in published literature(12-17, 23-27)(Supp. Tables 1, 2, and 3, sheet 3).

We observe that many nodes regulate more than one type of phenotypic output. For example, the MRPL9 protein is involved in both voltage generation and inheritance of the mitochondria(25). In the mouse brain, the lateral hypothalamus regulates feeding behavior and predator responses(33). In the worm, the RMGL and RMGR neurons function in egg-laying, feeding behavior, and CO_2_/O_2_ detection responses(36).

We rarely observe a situation in which all participating nodes in a phenotypic output are members of the same module. However, we find that nodes involved in the same phenotypic output tend to be connected to one another, and to form subnetworks with specific topological features. These ‘phenotypic subnetworks’ have high *Cg* values, as compared with both Erdos-Renyi and randomized subset controls, suggesting that they were generated by evolutionary selection. However, they often have *M* values that are less than those of one or both of the control subnetworks or of the communities within the network.

In Figure 3, we show diagrams for four phenotypic subnetworks for the yeast mitochondrial network. Figure 4 shows four phenotypic subnetworks, representing animal behaviors, for the mouse brain connectome, and Figure 5 shows four for the *C. elegans* connectome. (Supplementary Figures 5-7 show the same diagrams, but with the names of components included.) These phenotypic subnetworks are of very different sizes and span different numbers of sub-communities (modules). Figure 3 shows that mitochondrial inheritance involves nodes that are dispersed across 21 sub-communities, while mitophagy uses nodes that are largely, but not exclusively, within a single sub-community. Similarly, in Figure 5, we see that aggregation/dispersion behavior involves nodes within seven sub-communities, while feeding behavior involves a much smaller number of nodes that are restricted to four sub-communities. Despite these differences, all of the phenotypic subnetworks have significantly higher *Cg* values than the controls (Figures 3-5). Lists of all detected phenotypic subnetworks along with their topological parameters are provided in sheet 3 in Supp. Tables 1, 2, and 3.

**Figure 3.**
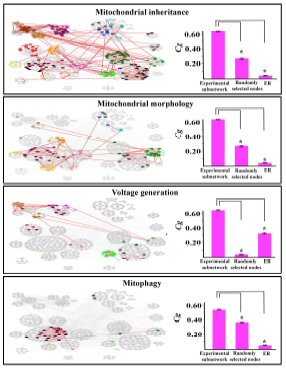
Example phenotypic subnetworks for the yeast mitochondrial network. Nodes (proteins) implicated in four network functions are superimposed in color on a gray background corresponding to the entire network diagram from Figure 1A. Connections (edges) within the phenotypic subnetworks are indicated by red lines. These diagrams show the phenotypic subnetworks of nodes implicated in mitochondrial inheritance (A), mitochondrial morphology (B), voltage generation (C), and mitophagy (D) in published papers (see references in main text). The color code indicating community and sub-community identity is the same as in Figure 1A. Identities of nodes are indicated in Supp. Figure 4. The histograms on the right side compare *Cg* for each phenotypic subnetwork to those of networks made by a random selection of the same number of nodes from the entire network, or by an ER model with the same number of nodes. Note that the *Cg* values for each of the phenotypic networks are much larger than those for the corresponding random or ER networks. P<0.0001 for all indicated comparisons (brackets).

**Figure 4.**
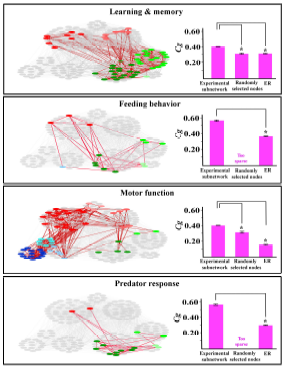
Example phenotypic subnetworks for the mesoscale connectome of the mouse brain. Nodes (brain regions) implicated in four network functions are superimposed in color on a gray background corresponding to the entire network diagram from Figure 1b. Connections (edges) within the phenotypic subnetworks are indicated by red lines. These diagrams show the phenotypic subnetworks of nodes implicated in learning and memory (A), feeding behavior (B), motor function (C), and predator responses (D) in published papers (see references in main text). The color code indicating community and sub-community identity is the same as in Figure 1B. Identities of nodes are indicated in Supp. Figure 5. The histograms on the right side compare *Cg* for each phenotypic subnetwork to those of networks made by a random selection of the same number of nodes from the entire network, or by an ER model with the same number of nodes. “Too sparse” indicates that the randomly selected network had too few connections to allow a calculation of its *Cg* value. Note that the *Cg* values for each of the phenotypic networks are much larger than those for the corresponding random or ER networks. *p*<0.0001 for all indicated comparisons (brackets).

**Figure 5.**
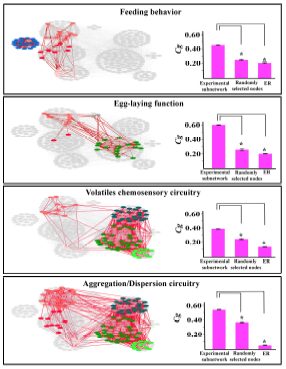
Example phenotypic subnetworks for the *C. elegans* connectome. Nodes (neurons) implicated in four network functions are superimposed in color on a gray background corresponding to the entire network diagram from Figure 1C. Connections (edges) within the phenotypic subnetworks are indicated by red lines. These diagrams show the phenotypic subnetworks of nodes implicated in feeding behavior (A), egg-laying (B), volatile chemosensation (C), and aggregation and dispersion (D) in published papers (see references in main text). The color code indicating community and sub-community identity is the same as in Figure 1C. Identities of nodes are indicated in Supp. Figure 6. The histograms on the right side compare *Cg* for each phenotypic subnetwork to those of networks made by a random selection of the same number of nodes from the entire network, or by an ER model with the same number of nodes. Note that the *Cg* values for each of the phenotypic networks are much larger than those for the corresponding random or ER networks. *p*<0.0001 for all indicated comparisons (brackets).

Taken together, the above results indicate that, although the walk-trap method is not sensitive to edge length, it is capable of capturing proximal modular structure in all of these biological networks. We also find that phenotypic outputs of the networks are products of interplay among nodes in different modules that are connected to form subnetworks with characteristic topological features, such as high *Cg* values relative to random networks of the same size. The existence of such characteristic features indicates that phenotypic subnetworks are products of selection, and may serve as statistical markers for physiologically relevant functions. However, it remains very challenging to identify phenotypic subnetworks *via* computational methods alone, and experimental approaches are still required for identification of the nodes within these subnetworks.

### A new computational model captures the global topological features of all three networks

The walk-trap algorithm analyses led to the conclusion that modular architecture is a key topological feature of all three networks. We find that large communities within the networks are well fit by a binomial/Poisson *P(k)* distribution, and that there is a higher density of edges within a community and a lower density between communities (Supp. Table 5). One way of generating this kind of topology is by using a stochastic block model (SBM)(37, 38), which consists of several interconnected Erdos-Renyi communities (blocks), with a higher probability of forming edges between nodes in the same block than between nodes that belong to different blocks (Figure 6A). We used an SBM model to generate network simulations with numbers of edges and nodes similar to those of the yeast mitochondrial, mouse brain mesoscale connectome, and *C. elegans* connectome networks. The SBM successfully captured modularity (*M*), and small-world properties (*L*) of all three real-world networks (Figures 6B, C, and D). However, although SBM produces networks with higher *Cg* than the ER, BA, and HRG algorithms, they are still too low for the SBM to describe the generation of the real networks. One possible reason for this relatively low *Cg* is the lack of a selection mechanism that reinforces important edges and removes less important ones. Also, there are substantial fitting errors, as measured by the SSE, for the *P(k)* distributions when SBM is compared to the real networks.

**Figure 6.**
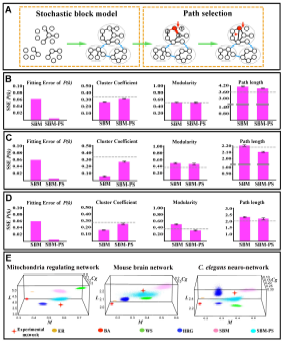
The SBM-PS model captures the key topological features of all three large biological networks. Cartoon representations of the SBM model (A, left panel), and the path selection algorithm in the SBM-PS model (A, right panel). In SBM, edges have a higher probability of forming between nodes in the same community (black lines) than between nodes in different communities (blue lines). For path selection, nodes and edges are ranked based on the number of paths passing through them. In the example, the numbers on three nodes and the thickness of the edges (red) that connect them to the red node indicate their ranking. Edge degradation rate is inversely proportional to edge ranking (red arrow indicates a removed edge). New edges (arrowhead) are preferentially attached to higher-ranking nodes. The SBM-PS model captures the major topological features of the yeast mitochondrial network (B), the mesoscale connectome of the mouse brain (C), and the *C. elegans* connectome (D). The SSE values for the divergence (fitting error) between the *P(k)* distributions of the experimental networks and those of the model networks are much lower for SBM-PS than for SBM, ER, WS, or BA. For *Cg*, only SBM-PS and WS generate high enough values. See Figure 1 for ER, WS, and BA histograms. The grey dashed line in the histograms represents the value of these topological parameter in the real network. (E) A 3-D representation of the distributions of possible values of *Cg*, *M*, and *L* for the various network models. Each colored region is a cloud of points, representing the possible values of these three parameters that can be produced by each of the models. The red crosses mark the actual values for the experimental networks. Note that the cross for the mouse brain connectome network is immediately adjacent to the turquoise cloud for the SBM-PS network, indicating that this model fits the experimental network excellently. For the yeast mitochondrial network and the *C. elegans* connectome network, the cross is much closer to the SBM-PS cloud than to any other model clouds. These data, together with the *P(k)* distribution histograms in panels B, C, and D, shows that SBM-PS is uniquely able to capture the key features of all three experimental networks.

At a local level, minimizing the shortest path between three connected nodes is best achieved when they are completely connected in a closed triangular formation, which is the basic unit used for *Cg* measurements. Our model randomly selects a node *v*_*0*_, then assigns weights to the edges connecting it to its neighboring nodes. The degree of a neighboring node *v*_*i*_ (that is, the number of other nodes to which it is connected) serves as a proxy for the weight of connected edges. Intuitively, edges that connect to nodes with high degree allow formation of more short paths than edges connected to nodes with low degree. We created a selection algorithm based on these principles that increases the rate of deletion of edges with low weights. We then coupled the SBM model, which preferentially forms edges between intra-community nodes, with the edge deletion algorithm. This composite model favors the preservation of edges that create closed triangles and thus increases *Cg* values locally. We refer to this new composite model as the Stochastic Block Model with Path Selection (SBM-PS)(Figure 6A).

To compare the SBM-PS model against the frequently used ER, BA, WS, HR, and SBM computational network models, we generated 20,000 simulated networks from each model with parameter values that gave as close to the same numbers of nodes and edges as the yeast mitochondrial, mouse brain, and *C. elegans* networks as possible. By measuring the quantitative properties of each simulated network, we are able to calculate the *P(k)* distributions, *Cg*, *M*, and *L* values for each model. Remarkably, we found that SBM-PS generates *Cg* values that are very close to those of the real networks, while preserving the *M* and *L* values generated by SBM (Figures 6B, C, and D; grey dotted lines indicate values for the real networks). Furthermore, the fitting error, as measured by the SSE, for the *P(k)* distributions was almost zero for SBM-PS (Figures 6B, C, and D), indicating that this model provides an excellent fit to the *P(k)* distributions of the real networks.

From these data, we are able to estimate empirical probabilities that the quantitative properties of the yeast mitochondrial, mouse brain, and *C. elegans* networks are as extreme as or more extreme than the simulated networks for each model. We combined these probabilities using Fisher’s and Stouffer’s method to produce a single combined probability for each network that reflects the probability that that network could be generated by each of the models(39-41). Lastly, we considered fitting a multivariate normal distribution to each of the four quantitative properties(39-41). While each of the resulting combined probabilities are low, it is clear that the probability that the three networks could have been generated by the SBM-PS is several orders of magnitude higher than for any competing model (Figure 6E, and Supp. Table 5).

We note that both the SBM-PS and Watts-Strogatz models can generate networks with high *M* and *Cg* values that are prerequisites for the formation of the experimentally observed phenotypic subnetworks. However, unlike the Watts-Strogatz model, the SBM-PS model can produce *P(k)* distributions that closely match those of the complete yeast mitochondrial, mouse brain connectome, and *C. elegans* connectome networks.

The SBM-PS model also produces ‘modular heterogeneity’ and ‘node heterogeneity’. These terms refer to the facts that modules of different sizes will have different edge formation probabilities, and nodes will have different wiring probabilities that are dependent on their neighbors and on the structure of the module to which they belong. Therefore, modules generated by the SBM-PS model are more representative of biological modules than modules generated by other computational algorithms, since each will have a unique architecture, structured by evolutionary selection, that facilitates the abilities of its nodes to execute their functions.

## Conclusions

There are three major findings that emerge from our analyses of the yeast mitochondrial protein network, the mouse brain mesoscale connectome network, and the *C. elegans* connectome network. First, despite the fact that the yeast mitochondrial network is a proteomic network of molecules within a cell, while the other two networks are anatomical networks of neurons or groups of neurons within an organism, all three networks have similar global topologies. Second, the functions of the networks, which we call phenotypic outputs, involve collections of nodes that span modular boundaries and form phenotypic subnetworks with particular topological properties. These results suggest that biological function is facilitated by specific kinds of interconnected networks. Third, we were able to develop a new model, SBM-PS, that captures the topological properties of these three very different networks more faithfully than other models. The communities within the network, which have binomial/Poisson-like *P(k)* distributions, can be modeled using other algorithms. We have previously shown that a fat-storage regulating network (which consists of ~1/8^th^ of the mitochondrial network) can be effectively simulated by the Watts-Strogatz network model(23). The fact that SBM-PS is the only model capable of recapitulating the topological attributes of the complete yeast mitochondrial network and of the mouse brain and *C. elegans* connectomes suggests that its path selection algorithm represents a plausible mechanism by which complex networks can be generated through evolution. In this mechanism, the process of selection increases the speed of communication within and between modules and communities in the network. Modules can represent different molecular complexes, as in case of the mitochondrial network, or different local neuronal circuits in the case of the mouse brain and *C. elegans* neural networks. By linking communities or modules that are already optimized to perform a given function into a single network, natural selection may be able to produce control systems that efficiently integrate many inputs and can generate diverse outputs.

## Acknowledgments

This work was supported by a grant from the NIH, R21NS083874, to K. Z., and by the Della Martin Foundation. We also like to acknowledge NVIDIA Corporation for generously donating the NVIDIA GTX980 Graphics Card used in this study

**Uncategorized References**

